# Mistimed feeding disrupts circadian rhythms of male mating behavior and female preovulatory LH surges in mice

**DOI:** 10.1101/2021.12.27.474267

**Authors:** Ayaka Kukino, Thijs J. Walbeek, Lori J. Sun, Alexander T. Watt, Jin Ho Park, Alexander S. Kauffman, Matthew P. Butler

## Abstract

In rodents, eating at atypical circadian times, such as during the biological rest phase when feeding is normally minimal, reduces fertility. Prior findings suggest this fertility impairment is due, at least in part, to reduced mating success. However, the physiological and behavioral mechanisms underlying this reproductive suppression are not known. In the present study, we tested the hypothesis that mistimed feeding-induced infertility is due to a disruption in the normal circadian timing of mating behavior and/or the generation of pre-ovulatory luteinizing hormone (LH) surges (estrogen positive feedback). In the first experiment, male+female mouse pairs, acclimated to be food restricted to either the light (mistimed feeding) or dark (control feeding) phase, were scored for mounting frequency and ejaculations over 96 hours. Male mounting behavior and ejaculations were distributed much more widely across the day in light-fed mice than in dark-fed controls and fewer light-fed males ejaculated. In the second experiment, the timing of the LH surge, a well characterized circadian event driven by estradiol (E2) and the SCN, was analyzed from serial blood samples taken from ovariectomized and E2-primed female mice that were light-, dark-, or ad-lib-fed. LH concentrations peaked 2h after lights-off in both dark-fed and ad-lib control females, as expected, but not in light-fed females. Instead, the normally clustered LH surges were distributed widely with high inter-mouse variability in the light-fed group. These data indicate that mistimed feeding disrupts the temporal control of the neural processes underlying both ovulation and mating behavior, contributing to subfertility.

## Introduction

Optimal timing of reproductive physiology and behavior is critical to ensure reproductive success (Morin et al., 1977; Goldman, 1999; Antle and Silver, 2016), and fertility can be compromised by circadian alterations in reproductive mechanisms. Circadian desynchrony occurs when endogenous circadian rhythms in the body and external environmental rhythms become uncoupled. Human epidemiological work and experiments in rodent models indicate that circadian desynchrony compromises reproductive health. Shift workers have an increased risk for poor reproductive outcomes (Mahoney, 2010), including irregular menstrual cycles (Lawson et al., 2011), endometriosis (Marino et al., 2008), infertility and miscarriage (Fernandez et al., 2016), and need for fertility treatment (Fernandez et al., 2020). In addition, shift workers that do get pregnant demonstrate increased incidence of preterm birth and low birth weight of their babies (Xu et al., 1994; Bodin et al., 1999; Zhu et al., 2004).

Circadian desynchrony also impairs fertility in rodents (Miller et al., 2004; Alvarez et al., 2008; Summa et al., 2012; Takasu et al., 2015; Schoeller et al., 2016; Swamy et al., 2018). Under normal conditions, both male and female rodents exhibit 24-h rhythms in reproduction that are entrained to the light-dark cycle (Snell et al., 1940; Beach and Levinson, 1949; Everett and Sawyer, 1950). Male sex behavior is rhythmic (Sodersten et al., 1981; Logan and Leavitt, 1992). These rhythms depend on the central clock in the hypothalamic suprachiasmatic nucleus (SCN), for after lesions of this area, males still express sexual behavior but the normal rhythms are lost (Eskes, 1984). Female sex behavior is also strongly rhythmic (Wang, 1924) and is controlled proximately by the SCN’s regulation of endocrine milieu (Harlan et al., 1980). Ovulation and the associated endocrine signals in particular are tightly controlled by the circadian clock (Everett and Sawyer, 1950). Ovulation is preceded by a circadian-timed surge of luteinizing hormone (LH) secretion (‘positive feedback’) that is triggered via a multisynaptic neural pathway that begins with the SCN (Khan and Kauffman, 2012), and includes hypothalamic and pituitary areas that are also rhythmic and contain circadian clocks (de la Iglesia et al., 2003; Resuehr et al., 2007; Robertson et al., 2009; Zhao and Kriegsfeld, 2009; Chassard et al., 2015; Gotlieb et al., 2019). Rhythms are thus an intimate component of reproduction and disruption of the underlying clocks contribute to infertility.

Meals have a strong effect on circadian clocks, and an under-appreciated risk in shift work is the robust change in food intake pattern towards more night-time eating (Shaw et al., 2019; Flanagan et al., 2020; Kosmadopoulos et al., 2020). This can be modeled in rodents by time restricted feeding (TRF). TRF in during the inactive phase is deleterious for optimal reproduction, reducing fertility by impairing successful mating (Swamy et al., 2018). TRF’s impact may be via internal misalignment of clocks, for extra-SCN circadian clocks in the brain and in peripheral organs entrain to the food cue while the SCN still entrains to the light-dark cycle (Damiola et al., 2000; Gooley et al., 2006; Verwey et al., 2009). At present, the specific physiological and behavioral mechanisms underlying the reproductive impairments induced by TRF remain poorly characterized. Given the importance of both properly timed circadian rhythms to normal reproduction and the ability of food timing to alter such rhythms, we hypothesized that mistimed feeding reduces fertility in two complementary manners, by desynchronizing both male sexual behavior and female’s preovulatory LH surge.

## Methods

Two experiments were conducted in mice to assess the effects of TRF on either male mating behavior (Experiment 1) and/or the timing of the E2-induced preovulatory LH surge in females (Experiment 2), as described below. All procedures were approved by the Institutional Animal Care and Use Committee of Oregon Health and Science University.

### Experiment 1 – Effects of TRF on mating behavior

#### Animals, housing, and cohorts

Experiment 1 was conducted with homozygous *mPer2^Luc^* male and female mice derived from founders originally purchased from Jackson Laboratories (B6.129S6-*Per2^tm1Jt^*/J, Strain Code: 006852) (Yoo et al., 2004). This mouse line enables circadian rhythm tracking of the bioluminescent fusion protein of the clock protein PERIOD2 and firefly luciferase (PER2∷LUC). *mPer2^Luc^* mice were bred and maintained 2-4/cage in Thoren ventilated cages (Model #1, 19.6 cm × 30.9 cm × 13.3 cm) with pelleted cellulose bedding (BioFresh Performance Bedding, ¼” pelleted cellulose, Absorption Corp, Ferndale, WA) and ad libitum water and food (LabDiet 5L0D). Cages were placed in light-tight cabinets (Phenome Technologies, Skokie, IL); temperature and humidity were 23°C and 45% and lights were on a 12h:12h schedule. To achieve sufficient sample size, this experiment studied mice across 3 cohorts of 12 pairs each, half fed only during the dark and half fed only during the light (n = 6 mice * 2 sexes * 2 feeding conditions * 3 cohorts = 72). Cohort 1 mice were from 4 litters and there were two sibling pairs. Cohorts 2 and 3 mice were from 7 and 5 litters, respectively, with no sibling crosses. There was no overlap in parents across the 3 cohorts.

#### Time restricted feeding (TRF)

For this experiment, during conditions of food restriction (i.e., TRF), food was made available to both males and females only during the dark or only during the light phase of the photocycle. Chow was dropped into the wire top hoppers at specified times by automatic feeders placed on top of the cage lid. Remaining food was removed manually 12 hours later and the feeders were reloaded. When male and female mice were paired for mating behavior tests, food was provided and removed manually, keeping the same 12 hour restricted pattern. Water was always available ad libitum.

#### PER2∷LUC in vivo imaging

Clock gene expression (PER2∷LUC bioluminescence) was analyzed in vivo in a subset of 4 males and 4 females from cohort 1 to confirm food entrainment of peripheral clocks. After 2 weeks of TRF, bioluminescence from the liver was measured 10h and 22h after lights-on (zeitgeber times (ZT) 10 and 22; lights-on at ZT0) as previously described (Xie et al., 2020). Briefly, mice were lightly anesthetized with isoflurane and injected s.c. with D-luciferin potassium salt (15 mg/kg, Promega, Madison, WI) in sterile phosphate-buffered saline. After shaving fur from over the liver and throat, images were captured 10 minutes after injection using an Electron Magnified (EM) CCD camera (ImageEM, Hamamatsu, Japan, controlled by Piper software version 2.6.89.18, Stanford Photonics, Stanford, CA) connected to an ONYX dark box (Stanford Photonics). Bioluminescence from circular regions of interest over each tissue was quantified using ImageJ (ImageJ, NIH) (Tahara et al., 2012; Swamy et al., 2018).

#### Locomotor activity

Passive infrared motion detectors mounted above cage lids were used to collect locomotor activity as counts per 10-minute bins (Telos Discovery Systems, West Lafayette, IN). Locomotor activity profiles were constructed using ClockLab (Actimetrics, Wilmette, IL). Locomotor activity patterns of males and females were analyzed for the last 7-days of TRF, ending 12 hours before they were paired.

#### Mating behavior

After 5-12 weeks of exposure to restricted feeding, mating behavior was assessed in three independent cohorts of 12 pairs each, evenly split between dark-fed and light-fed treatment groups (overall n = 18 mating pairs/TRF treatment). Males were introduced first to a new cage in order to acclimate; one hour later, the female was introduced (Park, 2011) and the pair remained together for 96 h. Within TRF treatment, half of the mice were paired at lights-off and half at lights-on to control for the time of pairing. Estrous cycle was not monitored, so the estrous stage on the day of pairing was not known; therefore, the trial lasted 96 hours to ensure each female had potential to enter a receptive phase.

Behavior was recorded continuously for 96 h by video in IR mode (FDR-AX33, Sony Corporation, Tokyo, Japan). Light was provided by white fluorescent illumination during the light phase and dim red light during the dark phase. To improve video capture during the night, the cages were indirectly illuminated with an IR lamp directed towards the opposite wall (DMetric IR Illuminator, 850nm). All mounting attempts, independent of female receptivity, and all ejaculations were scored manually from the video by one observer and verified by a second. Female sex behavior is typically quantified by a lordosis quotient (receptivity) and strongly depends on normal male mounting attempts. Rhythms, or TRF disruptions thereof, in male mounting behavior would therefore confound estimates of female sex behavior rhythms, so receptivity was not scored in this experiment. At lights-on and lights-off, food was manually provided or removed from the food hoppers to maintain the TRF treatments, and all females were inspected for the presence of a copulatory plug at these times. In cohort 1, pairs were tested in the standard home Thoren cages on pelleted cellulose bedding, with food in a Thoren wire lid. In cohorts 2 and 3, pairs were tested in custom acrylic cages on white Alpha-Dri bedding and food provided in 1.3cm hardware cloth hoppers on the side of the cage. In both cases, water was available from sipper-tube bottles through a water grommet in the side of the cage.

### Experiment 2 – TRF effects on the circadian LH surge (E2 positive feedback)

#### Animals and housing

C57BL/6 females (n=35), purchased from Charles River at 10 weeks of age, were housed 3-4/cage in light-tight cabinets at 23°C, with lights on from 0800-2000h PST (12:12 light cycle). Food and water were available ad libitum.

#### Experimental protocol

After 2 weeks of acclimation to the 12:12 light cycle, females were assigned to one of three feeding conditions balanced for body weight: dark-fed (n = 12), light-fed (n = 12), and ad-lib fed (n = 11). At this time, all females were housed 2/cage except for one cage of 3 in the ad-lib group. To control for potential effects of shifts in food timing, half the mice in each feeding group were maintained on the 0800-2000h schedule and the other half were transferred to another cabinet with a reverse 12:12 light cycle (lights on from 2000-0800h). Thus, half of the food-restricted mice received food during their original photophase and half during their original scotophase. Mice and their food were weighed weekly.

Beginning 5 weeks after initiation of the TRF feeding paradigms, all females were briefly handled daily to accustom them to the tail-tip serial bleeding procedure. At week 6, mice were lightly anesthetized with isoflurane and ovariectomized between ZT0 and ZT3. At the time of ovariectomy, females were also implanted with a s.c. capsule containing 0.75 μg 17β estradiol (E2, Steraloids, Newport, RI) in sesame oil (25 μg/mL, 1.2 cm of oil in Silastic tubing, 0.20cm ID, 0.32cm OD, Dow Corning, Midland, MI). This E2 implant is well-characterized to induce a circadian-timed pre-ovulatory surge of LH in the evening 2 days after implantation (Dror et al., 2013; Poling et al., 2017; Mohr et al., 2021). For the ovariectomy and implantation surgeries, all mice were treated with meloxicam 5 mg/kg p.o. for analgesia (Meloxicam Oral Suspension, MWI Animal Health, Boise, ID).

Beginning two days after E2 capsule implantation, small serial blood samples were collected from the tail-tip every 2 hours for 24 hours starting at ZT4. For each sample, 12 uL of whole blood was collected in 108 uL of assay buffer (0.2% bovine serum albumin, 0.05% Tween 20 in PBS, pH = 7.5) and then frozen at −80°C until processing. Blood LH concentration was measured in duplicate by an ultra-sensitive murine LH ELISA at the Ligand Assay and Analysis Core at the University of Virginia. This ultra-sensitive LH assay uses a capture monoclonal antibody (anti-bovine LH beta subunit, 518B7) and detection polyclonal antibody (rabbit LH antiserum, AFP240580Rb) with a functional sensitivity of 0.016 ng/ml. Intra-assay %CV was 2.5% and inter-assay %CV was 7.3%. A commonly used definition of an LH surge is a peak that is 2 standard deviations above the average morning concentration across mice (Dungan et al., 2007; Dror et al., 2013). Because the time of the LH surge was unknown for these mice, we adopted the same ≥2 SD convention, but calculated a mean basal concentration across all mice (n=35), where each mouse’s basal concentration was the average of its 8 lowest concentrations. Two mice had elevated levels across the 24h without any evident peak, so for all potential surges identified by the ≥2 SD method, we further required that the peak be at least 4 SDs above basal concentration within-mouse.

### Statistics

To analyze rhythms of male mounting attempts, counts were collected into 10 min bins, collapsed across days into a single 24 h interval, and then normalized by dividing by the total mount attempts within-mouse. The normalized mounting was analyzed by repeated measures cosinor analysis with independent variables of TRF condition and zeitgeber time (ZT, ZT0 defined by lights-on and ZT12 defined by lights off, and parametrized with 2 harmonics of 24h and 12h). Mouse was included as a random factor. This analysis was conducted for all mice, for those mice that ejaculated, and for those that did not. Cohort was included as a factor, but this was always non-significant and removed. Body weight, food consumption, and LH concentration were analyzed by repeated measures ANOVA, with follow-up pairwise comparisons by Tukey test. Proportions of males exhibiting mounting behavior or ejaculations and females with LH surges were analyzed by chi square. Circular statistics were performed to determine whether events across the day (sex behavior or LH surges) were clustered at a specific time (Rayleigh test), and confidence intervals for ejaculation time and LH surge time were calculated according to (Batschelet, 1981). To compare distributions of ejaculation and LH surge times, the data were assumed to fit von Mises distributions (circular analog of the normal distribution) and a parametric test for the concentration parameter was conducted (U2 test, Case II 0.45≤*r*≤0.70, Batschelet, 1981, p. 122; NCSS, 2021). Statistical tests were considered significant with an alpha of 0.05. Error bars illustrate standard errors of the mean unless otherwise noted.

## Results

### Experiment 1 – Effects of TRF on mating behavior

Within each cohort, ages were approximately balanced between the light- and dark-phase feeding (Supplemental Material, **Fig. S1**). The first cohort was around 39 weeks of age at pairing; the second and third cohorts were around 25 weeks old. In the first cohort, by measuring PER2∷LUC bioluminescence *in vivo* in a subset of mice, we confirmed that our TRF paradigm successfully entrained the liver in both light- and dark-fed males and females (Supplemental Material, **Fig. S2**).

Home cage locomotor activity measured before the mounting behavior tests was bimodally distributed, with peaks at approximately ZT0 and ZT12 (**Fig. 1**). Both groups were more active during the dark-phase, with activity gradually increasing over the first half of the dark phase (ZT12-18) in the dark-fed control group but decreasing over this interval in the light-fed group (repeated measures ANOVA, effect of food, F_1,34_=0.67, p=.42; time of day (3h bins), F_7,238_=38.5, p<.001; interaction, F_7,238_=15.0, p<.001). Light-fed males and females showed similar locomotor activity temporal profiles (**Fig. 1B**), but dark-fed females were less active than males (Light-Fed: effect of sex, F_1,16_=0.12, p=.73; time of day (3h bins), F_7,112_=23.5, p<.001; interaction, F_7,112_=1.25, p=.28).

**Fig. 1.**
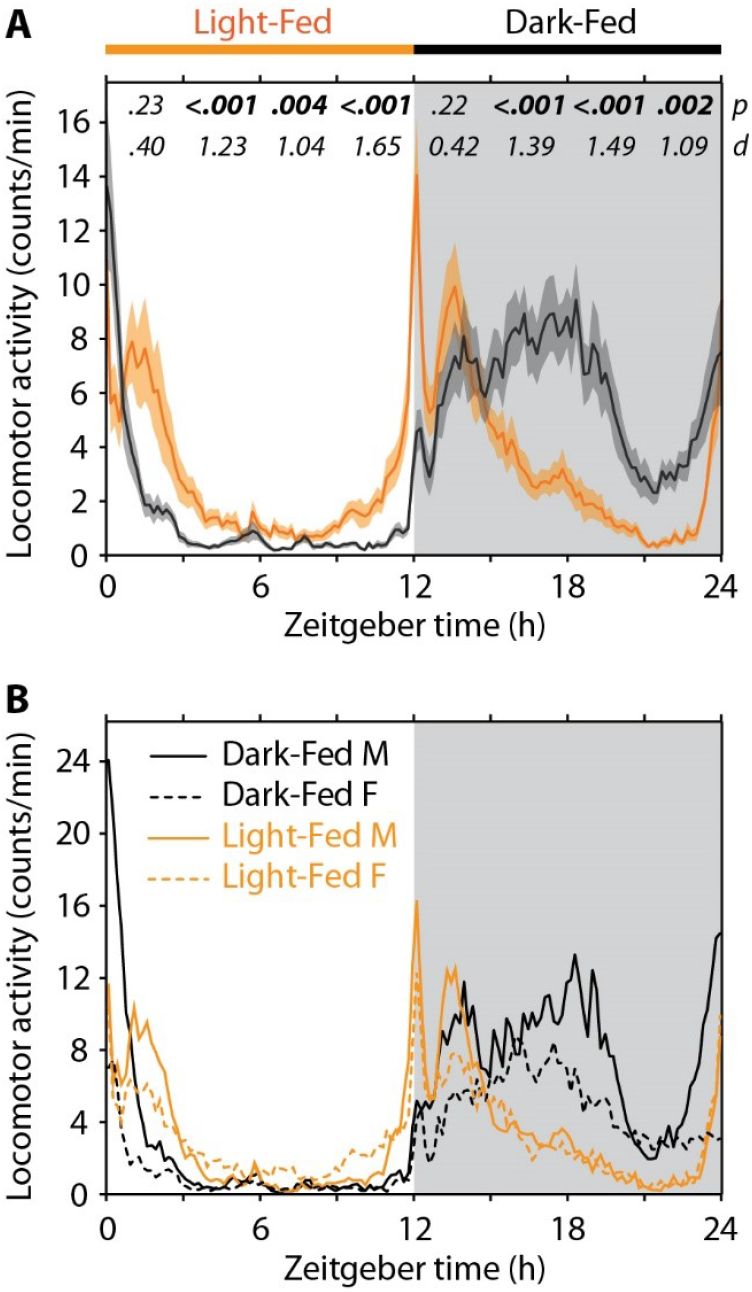
Locomotor activity during TRF. A. Spontaneous locomotor activity (mean and SE) in the week before mating behavior tests began. The dark period is shown in grey. Zeitgeber times 0 and 12 indicate lights-on and lights-off, respectively. Differences in activity level were assessed by t-test in 3h bins: p values and Cohen’s d are shown for each bin. N=18 cages per condition. B. There was little difference in the distribution of activity between males and females, though dark-fed females were less active than their male counterparts during the dark (Number of cages: 7 dark-fed male, 11 light-fed female, 8 light-fed male, 10 light-fed female).

Of the 36 mating pairs, one pair was separated in each TRF condition due to fighting, leaving 17 pairs per condition that completed the 96h behavior test. More of the males in the dark-fed control condition exhibited mounting behavior than in the light-fed group, but this did not reach statistical significance (**Fig. 2**. 15 versus 11, χ^2^(1,26)=2.6, p=.11). Light-restricted feeding significantly impaired mating outcome by halving the number of males that ejaculated (14 versus 7, χ^2^(1,21)=6.1, p=.014). Ejaculation was inferred from the presence of a copulatory plug in two dark-fed males; their ejaculation time was not measured due to loss of some video (see below).

**Fig. 2.**
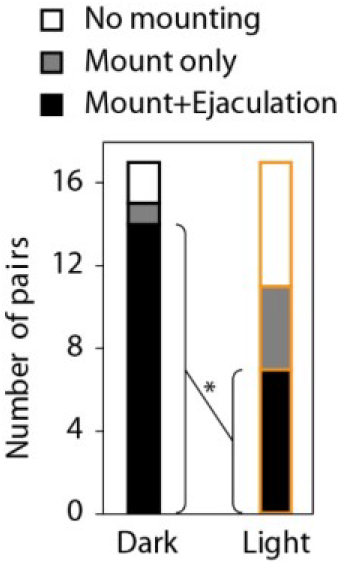
Mounting and ejaculation proportions. During the 96 h of pairing, there was a trend towards more dark-fed males exhibiting mounting behavior (p=.11), and significantly more of the dark-fed males ejaculated (*p=.014).

The temporal pattern of mounting behavior also differed between the two TRF groups (**Fig. 3)**. Dark-fed males displayed more mounting behavior during the late dark phase (ZT18-24), as expected since normal mating behavior typically occurs in the dark phase. By contrast, in the light-fed group, male mounting occurred more evenly across the day, with small peaks of mounting behavior in both the early light phase and early dark phase. There were significant interaction effects of TRF feeding condition with both the 24h and 12h time component in the analysis indicating that light-fed TRF treatment altered the normal temporal pattern of mounting behavior (**Fig. 3B**, F_1,4854_ = 14.1, p<.001 and F_1,4854_ = 38.0, p<.001, respectively; see Figure legend for complete statistics). Due to a technical error, video as lost for 24-36h in 6 dark-fed pairs. We therefore conducted two sensitivity analyses. First, we limited the data to only the last 72h of recording (3 pairs were still missing the first 12 h of this interval). Second, we repeated this 72h analysis without those 3 dark-fed mice without complete video records. In both cases, the conclusions were the same as the main analysis, with significant feeding condition × time interaction effects (Supplemental Material, **Fig. S3**).

**Fig. 3.**
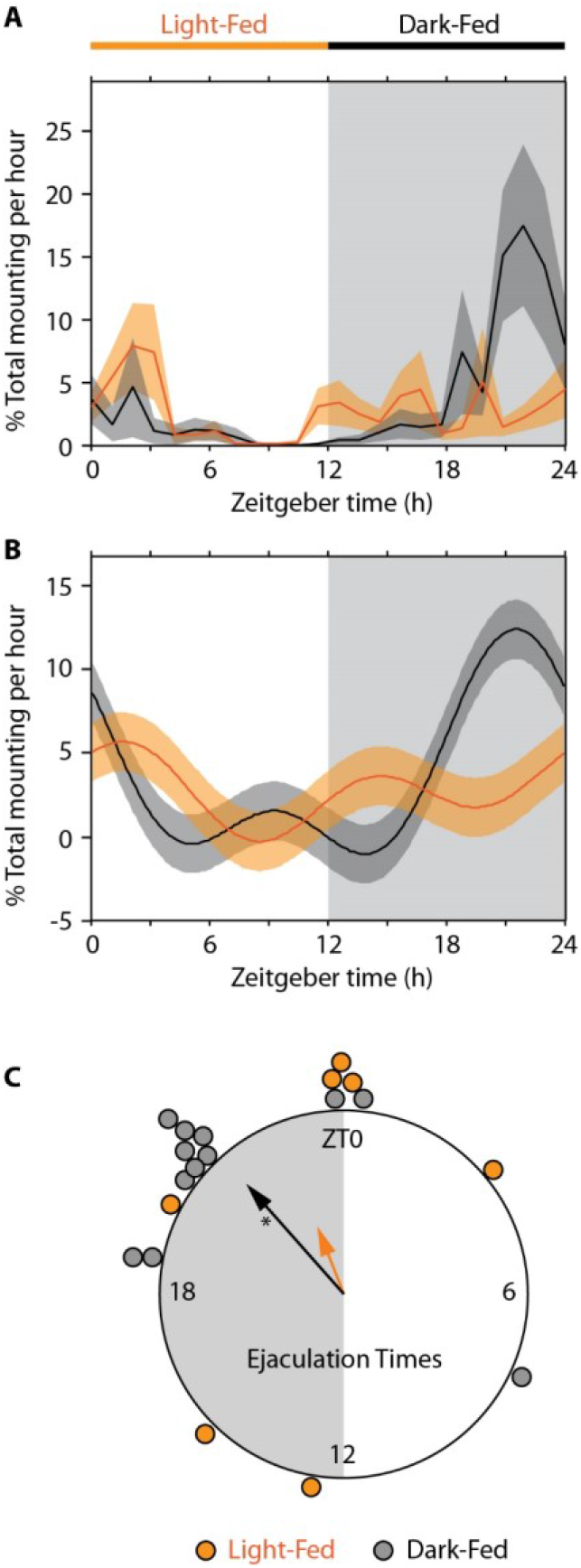
Mounting behavior and ejaculation timing. Normalized mounting behavior across all mice was collapsed over 24 h and averaged within condition (A, 1h bins). Dark-fed controls exhibited much more mounting in the late dark phase compared to light-fed males. Group differences were analyzed by a cosinor regression model (B, mean and 95% confidence interval). The best fit curve shows the same late night peak of mounting behavior in dark-fed mice and not light-fed TRF mice (effects of feeding condition, F1,32=2.67, p=.11; 24h rhythm F1,4854=53.7, p<.001; 12h rhythm, F1,4854=4.3, p=.038, feeding condition by 24h rhythm F1,4854=14.1, p<.001; feeding condition by 12h rhythm, F1,4854=38.0, p=<.001). C. Circular plot showing the time of ejaculations and the mean vector (arrow) for each group on the unit circle. *Mean vector length indicates significant clustering around a preferred time (Rayleigh test, p<.001).

Ejaculation times roughly followed the timing of mounting behavior (**Fig 3C**). The mean ejaculation time in dark-fed controls was ZT21.2 (95% confidence interval [19.4 – 23.0]), and was significantly clustered (*r*=0.76, Rayleigh test, p<.001, n=12). In contrast, though most ejaculations in light-fed TRF mice still occurred during the night, these were distributed widely (ZT22.5, 95%CI [16.5 – 4.5]) and there was no significant clustering in time (*r*=0.37, p=.39, n=7). When testing for a difference in how clustered the times were between conditions, the two groups did not differ (U2=1.57, analyzed as chi square with df=1, p=.21), but this may have been due in part to the small number of ejaculations in the light-fed group.

Mounting behavior rhythms were next re-analyzed in the subset of males that ejaculated or did not (**Fig. 4**). In males that ejaculated, similar to the whole group analysis above, there was a significant interaction effect of time and feeding condition such that dark fed males exhibited more mounting attempts during the late dark phase. In short, it was not the case that all ejaculating mice exhibited similar rhythms in mounting attempts. In contrast, in the males that did not ejaculate, mounting attempts were distributed much more evenly across the day, and though there was a small peak in the dark-fed mice, there was no significant TRF condition by time interaction effect (see Fig legend).

**Fig. 4.**
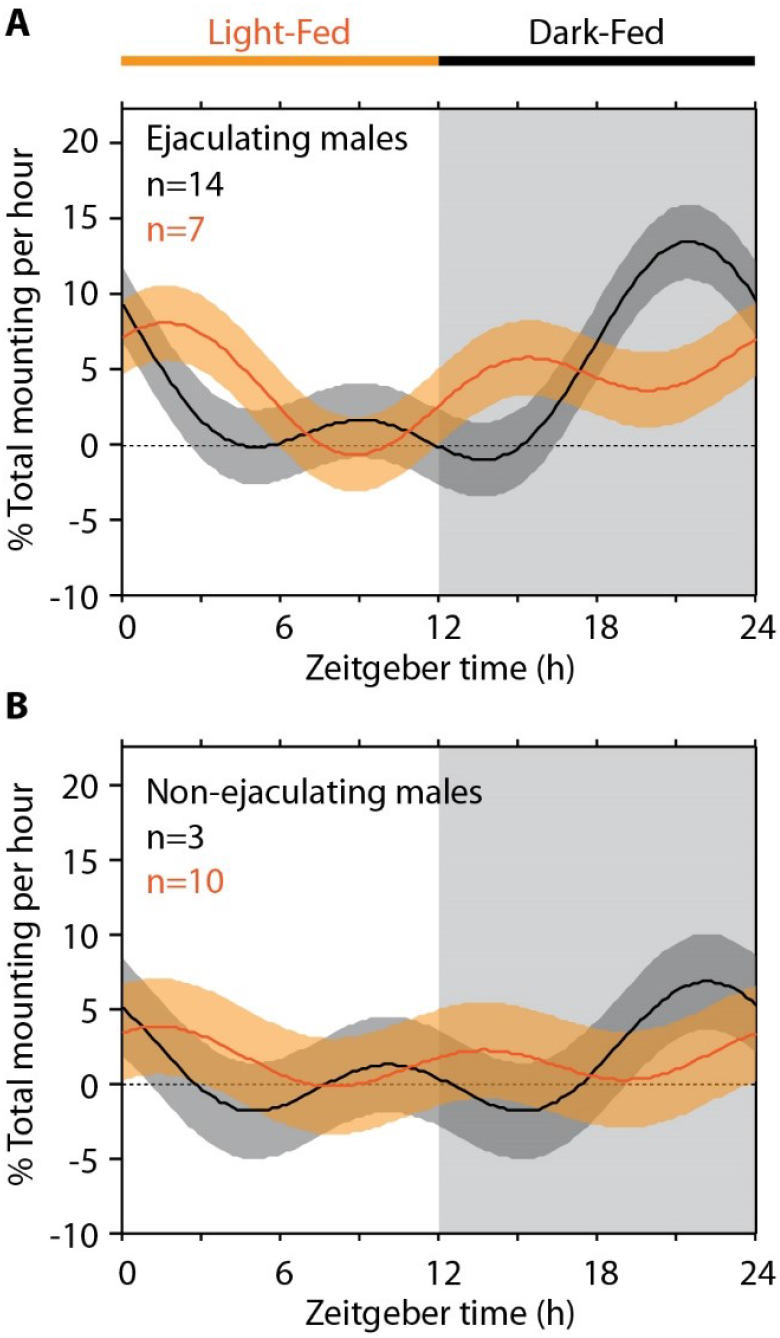
Mounting behavior timing (cosinor regression mean and 95% confidence interval) for males that ejaculated (A) or did not (B). Among those that ejaculated, mounting behavior in dark-fed mice peaked at the end of the night, while in light-fed mice, mounting was more widely distributed. Both groups had a trough in mounting attempts late in the light phase. Effects of feeding condition, F_1,19_=0.0, p=1.0; 24h rhythm F_1,3014_=37.6, p<.001; 12h rhythm, F_1,3014_=1.14, p=.29, feeding condition by 24h rhythm F_1, 3014_=5.06, p=.025; feeding condition by 12h rhythm, F_1, 3014_=26.4, p<.001. For males that did not ejaculate, there was less mounting behavior, with only a small late-night peak in the dark-fed and early morning peak in the light-fed during which the confidence interval did not overlap 0. In contrast to the full model (Fig 3) and the subset of males that ejaculated (A), there was no significant interaction of feeding condition and the 24h rhythm in the non-ejaculators. Effects of feeding condition, F_1,11_=0.04, p=0.85; 24h rhythm F_1,1851_=6.30, p=.012; 12h rhythm, F_1,1851_=3.66, p=.056, feeding condition by 24h rhythm F_1,1851_=1.51, p=.22; feeding condition by 12h rhythm, F_1,1851_=7.04, p=.0080.

### Experiment 2 – TRF effects on the circadian LH surge (E2 positive feedback)

Body weight and food consumption were tracked for the 5 weeks of TRF or ad lib feeding prior to experiments commencing (Supplemental Material, **Fig. S4**). Body weight dropped during the first week of TRF in the light-fed group, but thereafter all three groups gained weight similarly. There was no main effect of feeding condition (F_2,32_=2.6, p=.09), but significant effects of week (F_5,160_=108.9, p<.001) and week×condition (F_10,160_=5.2, p<.001). The light-fed group also ate slightly less than the two control groups, and all effects were significant (feeding condition, F_2,14_=7.5, p=.006; week, F_4,56_=13.3, p<.001; week×condition, F_8,56_=4.12, p<.001).

As expected, mean circulating blood LH concentrations exhibited a marked peak around ZT12-14 in both dark-fed and ad-lib control females, reflecting properly timed E2-induced LH surges that normally occur in the early evening, Conversely, the expected early evening LH surge was strikingly absent in the light-fed TRF females (**Fig. 5A**; main effect of zeitgeber time, F_11, 340_=2.80, p=.0016; effect of group, F_2,32_=2.26, p=.12, interaction F_22,340_=2.63, p<.001).

**Fig 5.**
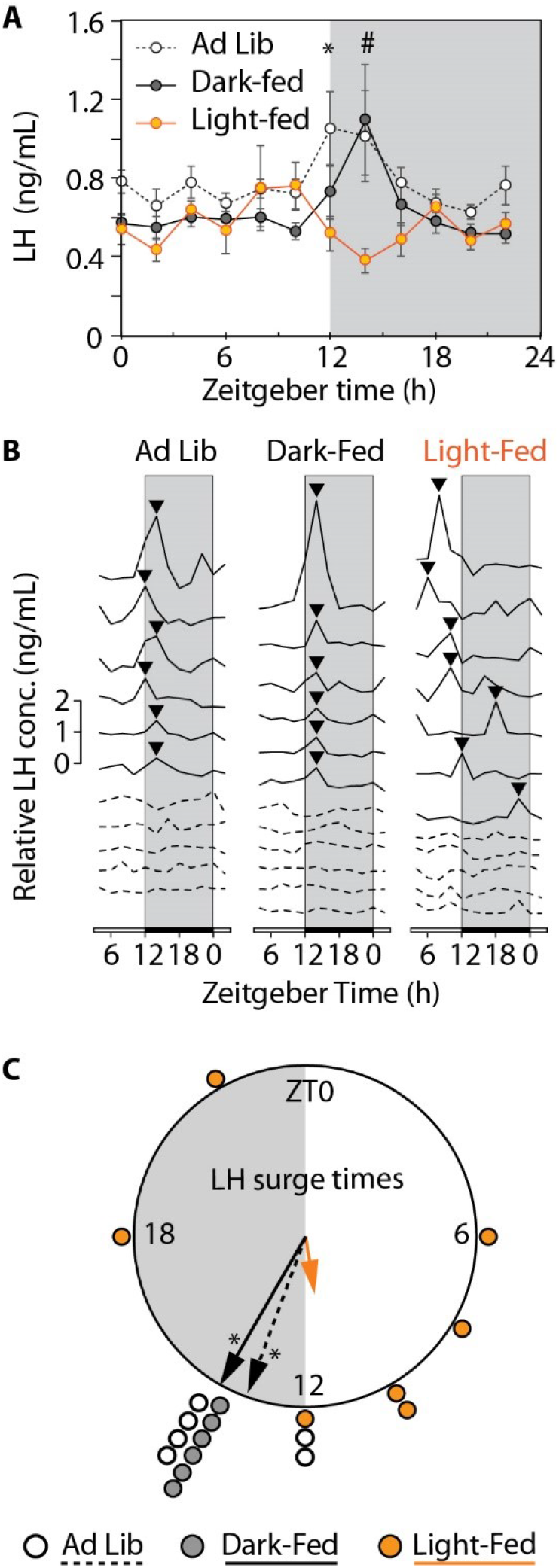
LH surge timing. **A**. E2-treated OVX mice in the ad lib and dark-fed control groups exhibited a rise in LH (mean and SE, n=11-12/group) at lights-off as expected. No such rise occurred in light-fed mice; their mean LH was significantly lower at ZT12 (†compared to ad lib mice, Tukey test, p<.05) and at ZT14 (#compared to both other groups, p<.05). Gray shading indicates darkness. **B**. Data from all individual mice show that the loss of the lights-off peak was due to desynchronization of the surges (▾) in the light-fed group. Overall, an LH surge was detected in about half of the mice in each group (solid lines compared to dotted lines). **C**. Circular plot showing the time of the LH surges and the mean vector on the unit circle. *Mean vector length indicates significant clustering around a preferred time in the ad lib and dark-fed groups (Rayleigh test, p<.001) but not in the light-fed group (p=.46).

LH surge temporal patterns were examined in individual mice to determine if the lack of an evening surge in the light-fed condition reflected the complete absence of LH surges throughout the day or desynchronized surges that occur at atypical circadian times rather than in the evening. Supporting the latter possibility, approximately half of the mice in each group exhibited a surge-like increase over a 2-4h period (no difference in proportion among groups, chi square test, χ^2^(df=2,35)=0.17, p=.92). In the ad lib and dark-fed control groups, these surge-like LH increases all occurred at ZT12-14, as expected (Rayleigh test for preferred time, *r*≥0.97, p<.001). In contrast, the surge-like increases in the light-fed females were desynchronized across mice, occurring at different times of the day (**Fig 5B**, *r*=0.34, p=.46). Within light-fed females showing such as surge, only 1 of 7 surges occurred at ZT12-14, significantly fewer than the 6 of 6 surges occurring at this circadian time in each of the dark-fed and ad lib-fed control groups (χ^2^(2,19)=15.0, p<.001). Surge times in light-fed mice were distributed significantly more widely than in both ad lib (U2=3.85, df=1, p=.0498) and dark-fed controls (U2=4.50, df=1, p=.034) (**Fig. 5C**).

## Discussion

Circadian disruptions, including those caused by mistimed food, are associated with a number of adverse health outcomes in animal models and humans, including impaired fertility, but the mechanistic underpinnings of such reproductive impairments are still not fully understood Here, in mice, we found that feeding restricted to the inactive phase of the light-dark cycle markedly disrupts the normal temporal organization of both mating behavior and the pre-ovulatory LH surge.

Disrupted reproductive rhythms suggest that food schedules impinge on the coordinated rhythmic brain circuits subserving the circadian timing of ovulation (Everett and Sawyer, 1950; Swann and Turek, 1985), including the circadian clock in the SCN, kisspeptin neurons in the AVPV, and GnRH neurons in the preoptic area (Khan and Kauffman, 2012; Kriegsfeld, 2013). The SCN transmits a daily signal to AVPV kisspeptin neurons via AVP release (Piet et al., 2015; Jamieson et al., 2021); when E2 is high, as during proestrus, AVPV kisspeptin cells respond robustly during the early evening and activate GnRH neurons that in turn elicit the LH surge (Robertson et al., 2009; Poling et al., 2017). All of these neural populations normally exhibit a circadian rhythm in activity and/or clock gene expression (de la Iglesia et al., 2003; Resuehr et al., 2007; Robertson et al., 2009; Zhao and Kriegsfeld, 2009; Chassard et al., 2015; Gotlieb et al., 2019). Thus, conditions or treatments that cause circadian desynchrony, such as TRF, may impede normal circadian activation of the neural mechanisms generating the LH surge. Give our present findings of mistimed LH surges in light-fed females, future studies are needed to determine if kisspeptin or GnRH neurons, or their upstream regulators, are affected by TRF, and how such effects might be induced. In some rodents, the LH surge also appears to determine the timing of female sexual receptivity (Fitzgerald and Zucker, 1976), though the neural circuits that govern sexual behavior are distinct from those controlling ovulation.

In prior studies of restricted food schedules, the SCN has been refractory to the entraining effects of food, such that misalignment between light and food cues is evident in misalignment between the SCN clock and clocks in peripheral tissues (Damiola et al., 2000; Stokkan et al., 2001). Importantly, TRF can also cause within-brain misalignment, by entraining extra-SCN brain areas, including the ventrolateral preoptic area (Neal-Perry et al., 2009) and dorsomedial hypothalamus (DMH) (Gooley et al., 2006; Verwey et al., 2009). Thus, one way TRF may interfere with reproductive function is by desynchronizing rhythms in kisspeptin or GnRH neurons, independent of the SCN. Relatedly, it is also possible that TRF enhances or activates upstream inhibitors of the LH surge generator circuitry, though this remains to be tested. This possibility is supported by observations that FOS protein (a marker of heightened neuronal activation) is induced in RFRP3 neurons of the DMH in anticipation of food during TRF (Acosta-Galvan et al., 2011) and that RFRP3, a neuropeptide that inhibits GnRH secretion, released at the wrong circadian time could impede the LH surge (Gotlieb et al., 2019). Moreover, it has been documented that RFRP-3 neuronal activation, as measured by FOS induction, is normally dampened at the time of the LH surge, supporting the possibility that the surge may be modulated by inhibition/disinhibition from RFRP3 signaling (Gibson et al., 2008).

Fertility may also be reduced by uncoupling the many behavioral rhythms that are normally synchronized for optimal reproduction: for example, uncoupling of rhythms in motivation for sex versus sexual behavior, as well as uncoupling of behavioral rhythms between males and females. In light-fed pairs, male mounting attempts and ejaculations were widely distributed, with no evidence for a preferred time. Ejaculations only occurred when females were receptive, but whether females were receptive at other times when the male was not attempting to mount is not known. The fact that the normally coordinated timing of mounting attempts, ejaculations, and LH surges were all disrupted in light-fed TRF conditions suggests that feeding impinges on normal physiology and behavior in both sexes. Nevertheless, males and females were acclimated to a common food schedule, so whether food schedules affect the reproductive behavior of one sex more than the other could not be determined.

Separate from restricted *timing* of feeding (TRF), caloric restriction has a well-documented inhibitory effect on female reproduction; underfeeding or reduced caloric intake can cause anestrus, and even mild caloric restriction can reduce aspects of mating behavior like partner preference (Bronson, 1989; Schneider et al., 2013). This may be mediated, in part, by a second population of kisspeptin neurons in the arcuate nucleus that indirectly regulate circulating estrous cyclicity and E2 concentrations via the direct control of GnRH pulse secretion (Clarkson et al., 2017; Wang and Moenter, 2020). Arcuate kisspeptin neurons are sensitive to metabolic signals and are inhibited in anorectic conditions (Padilla et al., 2017; Navarro, 2020). Nevertheless, in the current experiment, caloric intake was *not* restricted. Body weight and food consumption were tracked in the second experiment; light-fed mice ate slightly less and weighed slightly less than the two control groups. But all groups gained weight at similar rates throughout the experiment. Additionally, a change in E2 concentration is unlikely given that mice in the two groups do not differ in their cycling (Swamy et al., 2018). Regardless, in our LH surge study, all females were given exogenous E2 at proestrus levels to ensure all mice have similar E2; even in this scenario, light-fed females still exhibited alterations in their LH surge generation, indicating the reproductive problem is not due solely to insufficient E2 levels but rather includes neural impairments.

The circadian control of the LH surge may have evolved to ensure that ovulation occurs when sexual motivation and mating activity are high and the likelihood of encountering conspecifics is maximum (Morin et al., 1977; Simonneaux and Bahougne, 2015). In the field, where predation risk and resource availability are not uniform across the day, the circadian clock provides a competitive fitness advantage (DeCoursey et al., 1997; DeCoursey et al., 2000). Peripheral tissue clocks may provide further plasticity so that different physiological systems can be optimally timed to environmental cues (van der Veen et al., 2017). The circadian rhythms in the brain’s reproductive circuits may similarly play a role in fine tuning the occurrence of ovulation in variable conditions (Chappell et al., 2003; Zhao and Kriegsfeld, 2009; Khan and Kauffman, 2012). The ability to entrain reproductive timing to food availability may reflect an adaptation in environments more resource poor than the laboratory, even though TRF was still deleterious in this study’s lab setting. Conversely, and perhaps just as importantly, the reduced fertility itself may be the adaptation to avoid investment in reproduction when altered food timing might signal resource limitation.

Three minor limitations should be noted. First, because males and females were both on the same food schedules, we cannot identify any sex difference in the mating behavior response to TRF. Future studies could incorporate a design that exposes one sex to TRF while keeping the other on a normal feeding regimen. Second, female receptivity was not measured, though the wide distribution of ejaculation times in the light-fed TRF group suggests that receptivity is not tightly tied to the light-dark cycle, as is observed in normal mice. Because the males were also on TRF, female receptivity could only have been measured as a function of the male behavior. Third, these results are cross-sectional, so we cannot know if a mistimed LH surge would occur again at that same time (stably entrained to the same time within each mouse but desynchronized across mice) or whether the timing control is degraded such that a given mouse would also show cycle to cycle variability. The former result could indicate a compromise phase angle of entrainment to the competing cues of light and food. The latter result could indicate that the mice are failing to entrain stably to these cues. Because all mice were exposed to at least 5 weeks of TRF, they would not be expected to be still re-entraining to TRF.

Infertility and subfertility are important areas of public health that are sensitive to circadian disruption and shift work. The present conclusion that feeding restricted to occur solely during the biological rest phase disrupts the normal temporal patterns of the LH surge and mating behavior may have implications for how altered food timing typical of shift work or other circadian alterations may contribute to poor reproductive health. Reproductive health relies on precise coordination of timing in hypothalamic circuits – if disruptions thereof compromise ovulatory function and mating behavior, then timing meals to the active phase of the circadian cycle may prove therapeutic.

## Acknowledgements

The authors would like to thank Drs. Sato Honma, Wataru Nakamura, and Irv Zucker for helpful advice over the course of this project.

## Notes

<u>Funding:</u> This research is this publication was partially funded by the Oregon Institute of Occupational Health Sciences Innovation Project Funds and by NIH grant R01 NS102962 (MPB). ASK is supported by NIH grant R01 HD090161.

### Competing Interest Statement

The authors have declared no competing interest.

