## Supplemental Material for "Mistimed feeding disrupts circadian rhythms of male mating behavior and female preovulatory LH surges in mice"

**Figure S1.** Age of mice (mean, SE) when paired for behavioral testing. t-test, **p<.01, ***p<.001.

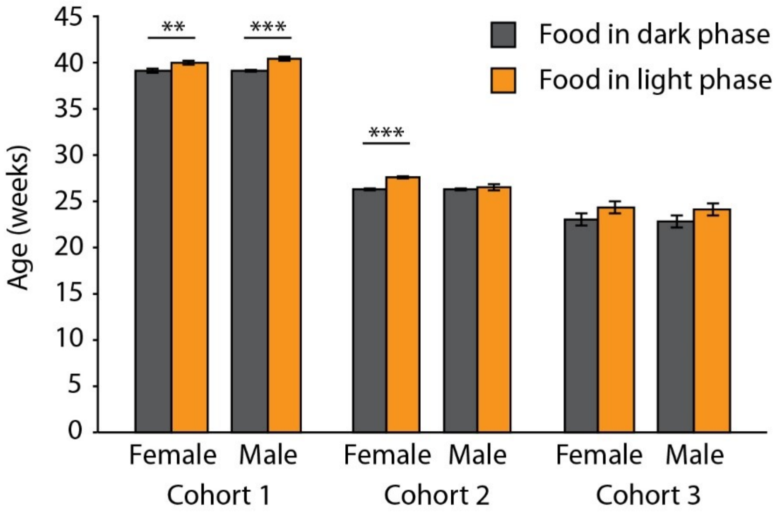


**Figure S2.** Liver PER2::LUC bioluminescence (mean, SE) was measured in vivo twice in a day from a subset of cohort 1 mice to verify that the food schedule had entrained peripheral organs, independent of the light-dark cycle. Bioluminescence was normalized to 100% within-mouse.

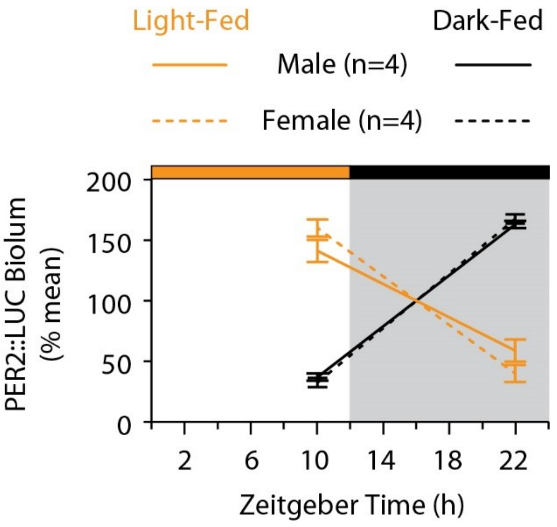

**Supplementary Material**

**Figure S3.** Sensitivity analysis shows that removing the first 24h of data or removing the first 24h of data and excluding the three mice without the full 72 h of data did not appreciably affect the finding that dark-fed mice have a peak of mounting behavior in the late night that is absent in day-fed mice. The interaction term of food condition by the 24 h circadian component indicated the same group difference in all three analyses.

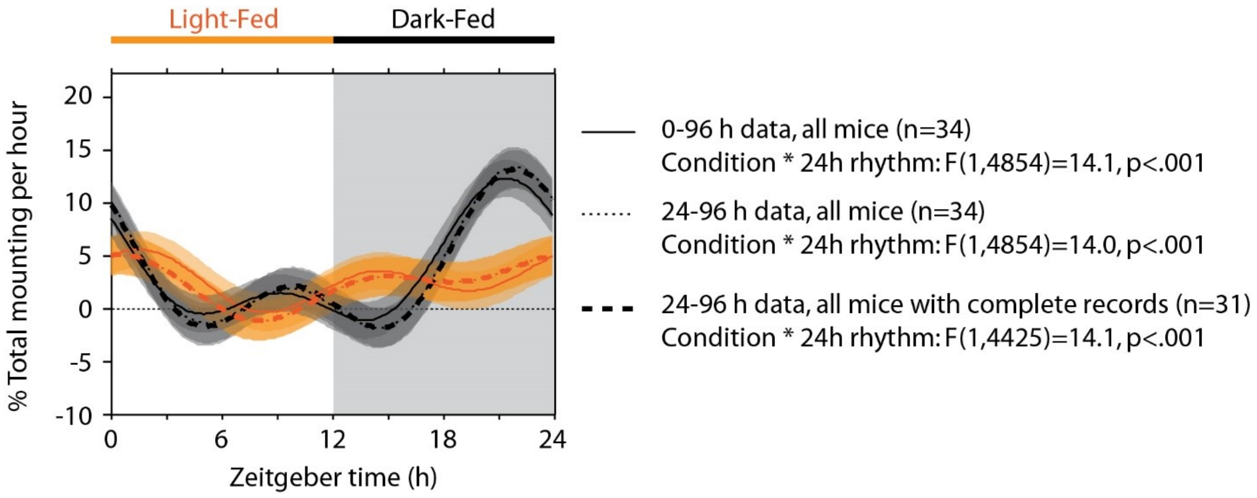


**Figure S4.** Body weight and food consumption (mean+/-SE) in females during acclimation in the LH study. Body weights were measured once a week and are the average of the body weight at ZT0 and ZT12. Females were group-housed. Therefore, food consumption was measured per cage and then expressed in g/day/mouse. Baseline food consumption was not measured. Week 0 represents the day before food restriction and light-cycle shifts were implemented. Significant pairwise differences are indicated (Tukey test, p<.05, indicated by: *Light-fed vs. ad lib; **Light-fed vs. both other groups; #Dark-fed vs. both other groups).

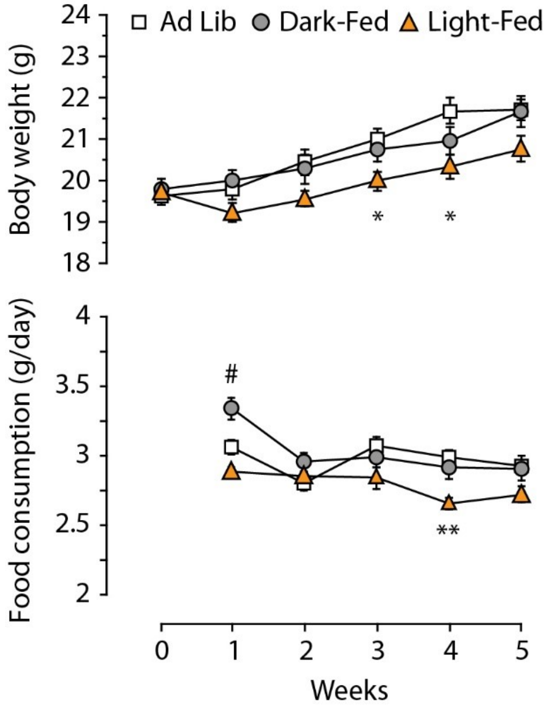
